# The variant Senescence-Associated-Secretory-Phenotype induced by centrosome amplification constitutes a pathway that activates Hypoxia-Inducible-Factor-1α

**DOI:** 10.1101/2020.12.08.415752

**Authors:** Selwin K. Wu, Juliana Ariffin, Remigio Picone

## Abstract

The Senescence-Associated-Secretory Phenotype (SASP) promote paracrine invasion however may also suppress tumour growth, thus generating complex phenotypic outcomes. Although centrosome amplification can induce proliferation arrest, the subsequent fate of cells with centrosome amplification remains elusive. Here, we report that centrosome amplification induces a variant SASP, constituting a pathway activating paracrine invasion. The centrosome amplification SASP is non-canonical as it lacks detectable DNA damage or prominent NF-κB activation. Instead, involving Rac activation and reactive oxygen species production. Consequently, inducing hypoxia-inducible factor 1α and associated genes, which includes pro-migratory factors such as ANGPTL4. Since senescent cells also have poor fitness, it is tempting to speculate that centrosome amplification induced SASP is one explanation for why extra centrosomes promote malignancy in some experimental models but are neutral or inhibitory in others.

## INTRODUCTION

Cellular senescence acts as a potent barrier for tumour development by promoting immune clearance of senescent cells and is triggered by anti-proliferation stresses such as constitutively active oncogene induction in untransformed cells^1^. The Senescences-Associated Secretory Phenotype (SASP) is a defining feature of senescent cells and is regulated at the transcriptional level by NFkB as a consequence of persistent DNA-damage^2, 3^.

Although centrosome amplification induced cell proliferation arrest^4, 5^, whether centrosome amplification induces SASP remains elusive. Centrosomes promote the assembly and organization of microtubules, which supports many cellular processes such as cell division, adhesion^6^ and migration^7^. In most normal cells, centrosomes are duplicated once during each cell division such that G1 phase cells typically have one centrosome while mitotic cells have two. Notably, the acquisition of supernumerary centrosomes is deleterious to untransformed cells. Untransformed cells with centrosome amplification activates signalling pathways that promote p53 stabilisation leading to cell proliferation arrest^4^. Perhaps counterintuitively, cancer cells commonly exhibit increased numbers of centrosomes. Thus, centrosome amplification often promotes spontaneous tumourigenesis in mice models when p53 is either deleted or becomes suppressed^9, 10^.

The senescence program is variably characterised by several non-exclusive markers, including constitutive DNA damage response signalling, senescence-associated β-galactosidase (SA-βgal) activity, increased expression of the cyclin-dependent kinase (CDK) inhibitor p21CIP1 (CDKN1A) and increased secretion of many extracellular factors through the SASP program^11^. Many senescence-associated markers result from altered transcription, but the senescent phenotype is variable. This variability is attributed to tissue and cell-type differences, differences in the senescence trigger^11^ and the dynamic nature of gene expression in senescent cells^12–14^. Despite variabilities in the senescent phenotype, a conserved transcriptome signature can be used as a useful marker to consistently identify senescent cells^15^. Here we report a pathway for the centrosome amplification induced a variant SASP to activate the Hypoxia-Inducible Factor 1-α in epithelial cells.

## RESULTS & DISCUSSION

### Centrosome amplification induced a variant senescence-associated secretory phenotype (SASP) independent of DNA damage

Supernumerary centrosomes were generated in cells by overexpressing Polo-like kinase 4 (PLK4), the master regulatory kinase for centrosome duplication ^16, 17^. PLK4 was induced transiently, and subsequent analysis was performed at time points where increased PLK4 mRNA was no longer detectable^18^. The majority of MCF10A cells (approximately 80%) contains extra centrosomes after 48 hours of PLK4 induction (Supplementary Fig. 1a). In most experiments described below, the transient overexpression of a truncated PLK4 mutant (Control 608) with kinase activity but does not localise to the centrosomes^19^ was used as the negative control.

SASP often induce paracrine invasion^20^. To determine whether cells with centrosome amplification could induce paracrine invasion, we used an established invasion assay with matrigel coated transwell^21^. MCF10A cells with or without centrosome amplification were introduced into the bottom chamber of transwells separated from the top chamber by a matrigel insert. Invasion of MDA-MB468 cancer cells across the matrigel and filter barrier (Fig. 1a) was then measured. Strikingly, centrosome amplification induced an ~2.5-fold increase in invading MDA-MB468 relative to the controls lacking centrosome amplification (Fig. 1a,b,c). Thus, soluble factors from cells with centrosome amplification can induce cell motility of nearby invasive cells.

**Figure 1.**
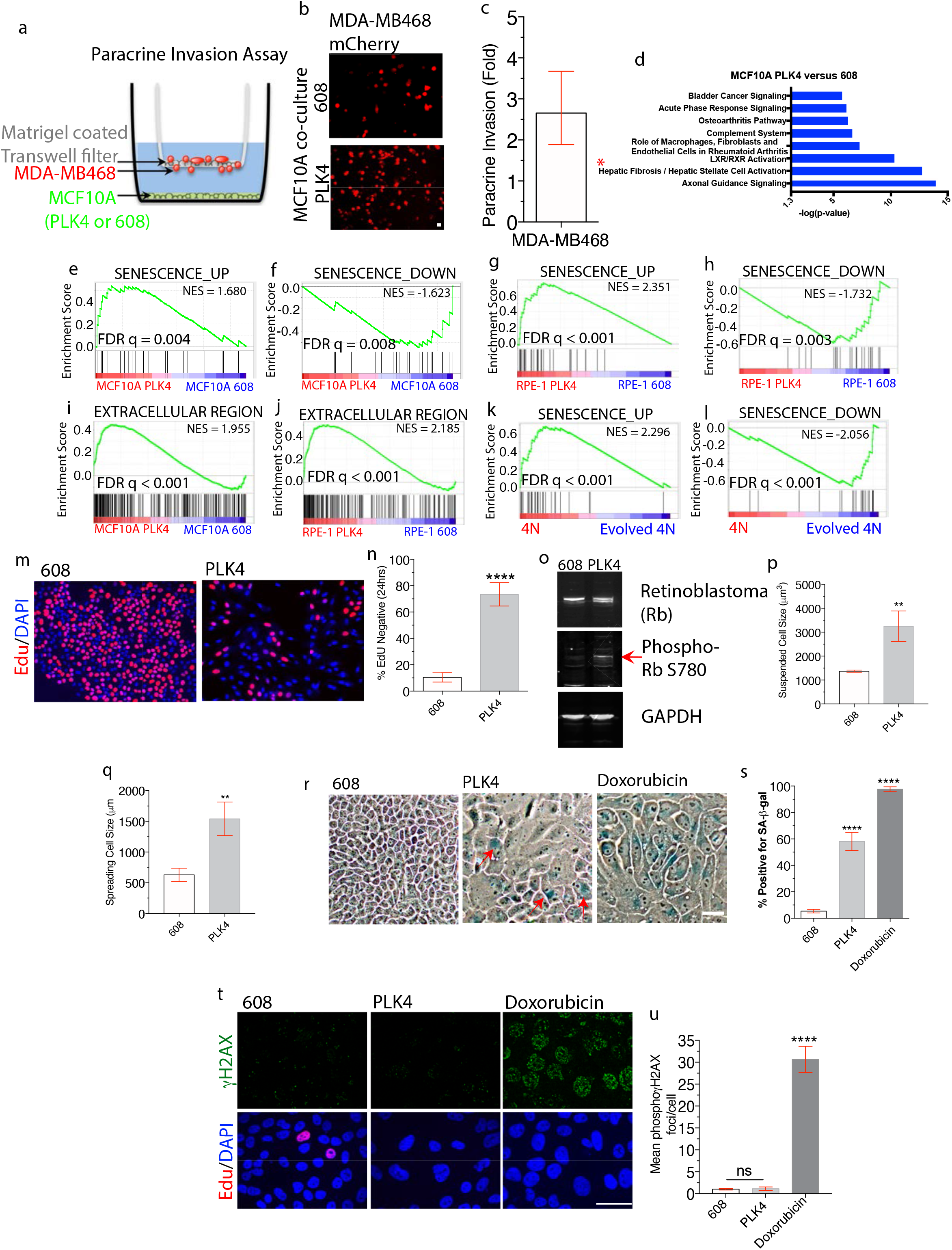
Centrosome amplification triggers a variant senescence-associated secretory phenotype (SASP). (a) Experimental scheme for the matrigel-coated transwell invasion experiment. Transwell inserts physically separates mCherry MDA-MB468 cells (top chamber) from the indicated MCF10A cells (bottom chamber). Paracrine invasion is scored from the numbers of mCherry MDA-MB468 cells that migrate to the bottom chamber. (b,c) MCF10A cells with centrosome amplification promote the invasion of MDA-MB468 mCherry cells. Representative images (b) and fold increase (c) of mCherry MDA-MB468 cells that crossed the matrigel-coated transwell upon co-cultured with the indicated MCF10A cells. (d) Centrosome amplification alters expression of genes related to cell motility and senescence. Ingenuity Pathway Analysis (IPA) of gene expression changes in MCF10A cells with centrosome amplification relative to controls revealing the top pathways altered by centrosome amplification. *Hepatic Fibrosis/ Hepatic Stellate Cell Activation is a senescence regulated process^22^. (e-h) Induction of senescence-related genes expression in cells with centrosome amplification. Gene set enrichment analysis (GSEA) revealed strong enrichment of genes up-regulated (e,g) or down-regulated (f,h) with senescence in MCF10A cells with centrosome amplification (e,f) and RPE-1 cells with centrosome amplification (g,h) relative to controls. NES: normalised enrichment score; FDR: false discovery rate. (i,j) Induction of secreted protein expression in cells with centrosome amplification. Gene set enrichment analysis (GSEA) revealed enrichment of genes annotated to be the extracellular region in MCF10A cells with centrosome amplification (i) or RPE-1 cells with centrosome amplification (j). NES: normalised enrichment score; FDR: false discovery rate. (k,l) Induction of senescence-related genes expression in cells with centrosome amplification. Gene set enrichment analysis (GSEA) revealed strong enrichment of genes up-regulated (k) or down-regulated (l) with senescence in RPE-1 tetraploids relative to “evolved tetraploids”. NES: normalised enrichment score; FDR: false discovery rate. (m,n) Proliferation arrest after centrosome amplification. Representative images (i) and quantification (j) of cells that cycled through S-phase (24 hr. EdU-label, red) and DAPI (blue) in MCF10A cells with (PLK4) and without (608) centrosome amplification. (o) Centrosome amplification induces retinoblastoma protein phosphorylation. Rb, Phospho Rb S780 and GAPDH (loading control) immunoblots of lysates from MCF10A cells with and without centrosome amplification. (p,q) Centrosome amplification increases cell size. Suspended (trypsinised) (j) and adherent (k) cell size measured for MCF10A cells with or without centrosome amplification. (r,s) Increased SA-β-Gal staining in cells with centrosome amplification. Representative images (n) and quantification (o) of SA-β-Gal (blue) staining of MCF10A cells with and without centrosome amplification or positive control with doxorubicin treatment. (t,u) Centrosome amplification does not induce DNA-damage. Representative images (p) and quantification (q) of γH2AX foci (green) in MCF10A cells with (PLK4) and without (608) centrosome amplification as compared to DNA damage from doxorubicin treatment. Cells in S-phase are labelled with Edu(red) pulse and excluded from the quantification. All data are means + s.e.m. from n=3 independent experiments,**P < 0.01,****P <0.0001; analysed with Student’s t–test except m which was with one-way ANOVA, Tukey’s multiple comparison test. Scale bars, 50 μm.

Next, we used RNA sequencing to identify SASP factors induced by centrosome amplification. Transient PLK4 induction was used to induce centrosome amplification in MCF10A cells and RPE-1 cells, resulting in ~86% and ~91% of cells respectively with centrosome amplification (Supplementary Fig.1a, b). The transcriptomes of cells with (PLK4) or without (Control 608) centrosome amplification were compared seven days post-induction. Among 1987 genes with significantly altered expression in MCF10A cells (fold change (FC) > ± 2, q-value < 0.05), 1285 were upregulated after centrosome amplification. For RPE-1 cells, 49 genes were significantly upregulated in cells with extra centrosomes. In both cell types after centrosome amplification, Ingenuity Pathway Analysis (IPA) revealed that the differentially expressed genes (fold change > ± 2, q-value <0.05) were enriched for terms associated with inflammation and the senescence-associated secretory phenotype (SASP)^22–24^ (Fig. 1d, Supplementary Fig. 1c).

Subsequently, we queried the transcriptomes of cells with (PLK4) or without (Control 608) centrosome amplification for gene expression signatures of senescence and extracellular protein induction. Despite variabilities in the senescent phenotype, a transcriptome signature have been identified to consistently identify senescent cells^15^. Thus, we employ this core senescent gene set to confirm the senescent identity of the whole-transcriptome-datasets analysed here. Gene set enrichment analysis (GSEA)^25^ in both MCF10A and RPE-1 cells revealed a strong association between centrosome amplification and senescence (FDR q=0.004, Fig 1e, FDR q=0.008, 1f, FDR q<0.001, 1g, FDR q=0.003, 1h) and extracellular region proteins(FDR q<0.001, Fig. 1i, FDR q<0.001, 1j). The top 50 genes in the “extracellular region” GSEA leading-edge subset from both cell types contain a wide variety of secreted factors, including extracellular matrix proteins, angiogenic factors, cytokines, and pro-invasive factors including the SASP factor IL8 (Supplementary Fig 1d,e,).

As an orthogonal approach to examine the effects of centrosome amplification, we employed a PLK4 expression-independent approach to induce extra centrosomes. We compared the transcriptomes of newly-generated tetraploid cells from dicytochalasin B induced cytokinesis failure with doubled centrosome content to controls. Both parental diploid cells and “evolved” tetraploid cells that had spontaneously lost their extra centrosomes are used as controls ^4, 26^. By IPA or GSEA with both data sets, the upregulated genes in newly generated tetraploids were enriched for secreted extracellular region proteins (FDR q<0.001, Supplementary Fig 1f,g), inflammatory and senescence-associated pathway terms (FDR q<0.001, Fig 1k,l and Supplementary Fig. 1h-k). The senescence-associated pathway terms are in common with what we observed in MCF10A cells exposed to 12Gy of γ-irradiation (Supplementary Fig. 1l). Our data show that the loss of extra centrosomes in tetraploids is sufficient to down-regulate senescence-related gene expression changes. Thus, orthogonal experimental approaches suggest that centrosome amplification induces senescence and a senescence-associated secretome.

Senescence is a complex and multifaceted phenomenon. Under current convention, cells are typically defined as senescent if they display proliferation arrest and at least two canonical senescence features^28, 29^. In addition to proliferation arrest (Fig 1m,n) and the SASP-like gene expression pattern, centrosome amplification induced other features of senescence. Consistent with a p53-dependent proliferation block, retinoblastoma phosphorylation was increased in cells with centrosome amplification (Fig. 1o). Also consistent with entry into senescence^30^, centrosome amplification induced increased cell size as assayed by measuring the volume of cells in suspension following trypsinisation (~2 fold increase, Fig. 1p) or by measurement of adherent cell area (~2.4 fold increase, Fig. 1q). The induction of senescence-associated beta-galactosidase activity (~60% positive cells, Fig. 1r,s) was also observed in cells with extra centrosomes, albeit at a lower intensity and frequency than was observed after extensive DNA damage. Persistent DNA damage response leading to DNA damage foci marked by γH2AX, 53BP1, or ATM (Ataxia-Telangiectasia Mutated)^2, 31^ is required for a robust SASP. However, multiple studies demonstrate that centrosome amplification does not induce detectable DNA damage in the nucleus^4, 10, 32^. Strikingly, even ~8 days after the induction of extra centrosomes, when many features of senescence are apparent, we observed no significant increase in the percentage of γH2AX-positive cells in the nucleus (Fig. 1t,u). Therefore, we reasoned that the weaker senescence detected by senescence-associated beta-galactosidase activity is due to a lack of extensive persistent DNA damage essential for a robust senescence phenotype.

### Centrosome amplification promotes the induction of ANGPTL4

We next sought to identify strongly differentially expressed genes common to all conditions, including cytokinesis failure, where centrosome amplification was induced. From our gene expression profiles in different cell lines, 7 genes were consistently upregulated in all experimental conditions (Fig 2a, Supplementary Fig 2a and Supplementary Table 1). These included the cyclin-dependent kinase inhibitor p21 (Supplementary Fig 2b), which is induced in all known examples of senescence, and IGFBP3, a known inducer of senescence^33^(Fig 2a,b). Among all conditions and cell types, ANGPTL4 was the most strikingly induced gene after centrosome amplification (up to ~15-fold induction in MCF10A cells, p<0.001 and approximately 5-fold in RPE-1 cells, Fig 2b,c). The induction of ANGPTL4 protein was confirmed using an enzyme-linked immunosorbent assay (ELISA) (Fig 2d). Importantly, inhibition of ANGPTL4 with a blocking-antibody or by CRISPR-mediated gene disruption significantly reduced paracrine migration stimulated by MCF10A cells with extra centrosomes (Fig 2e,f and Supplementary Fig 2c-f). These findings are notable because ANGPTL4 is a well-established pro-invasive factor that activates Rac signalling and disrupts cell-cell contacts^34–37^. As expected, the increased expression of ANGPTL4 is a common feature of SASP^38^.

**Figure 2.**
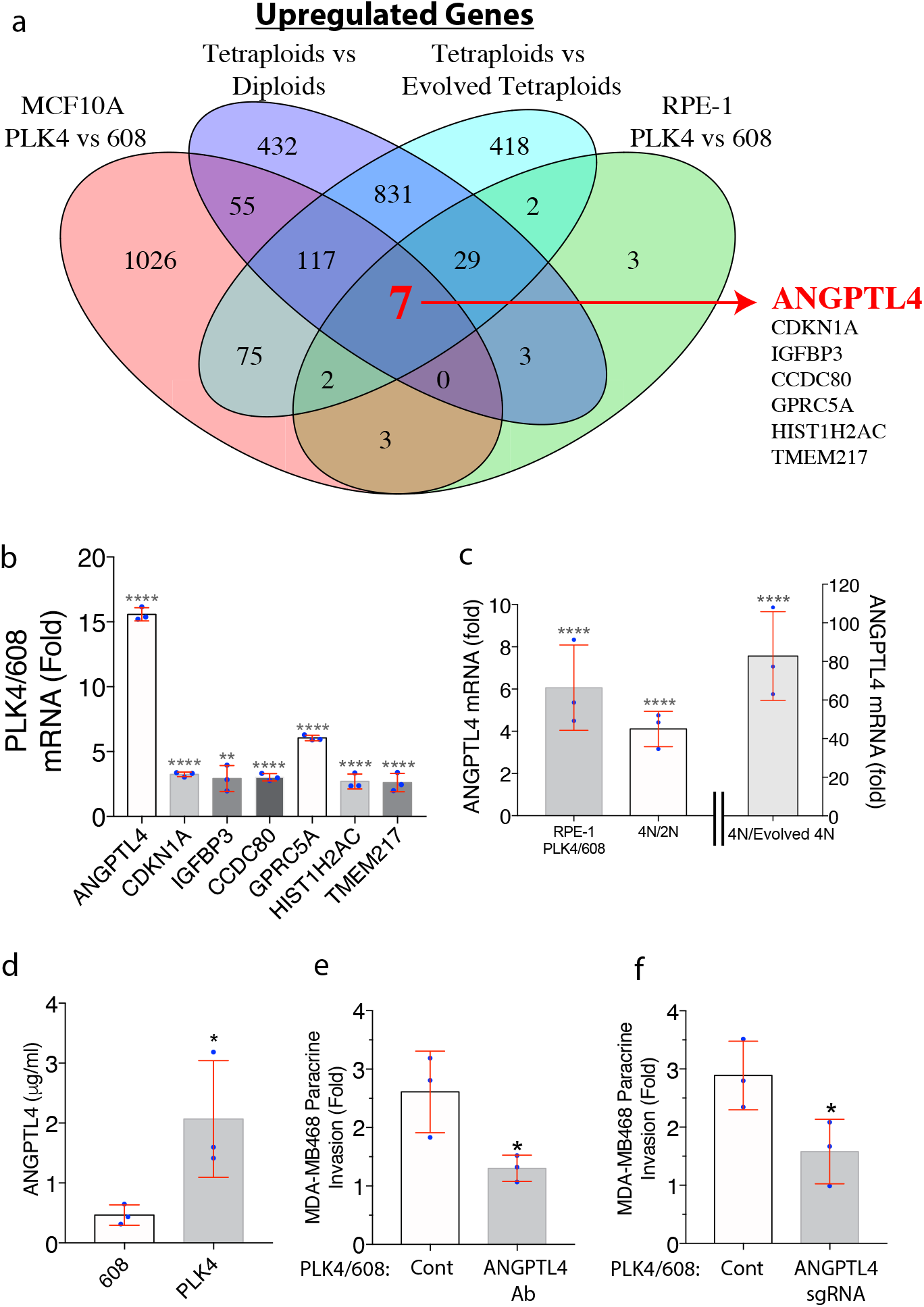
Centrosome amplification induced ANGPTL4. (a) Seven genes are upregulated by centrosome amplification from all experimental conditions, in two cell types. Venn diagram for the indicated comparisons showing the overlap of upregulated genes from RNAseq (2 fold change, p(adjusted)<0.05). (b) Among the 7 commonly upregulated genes, ANGPTL4 expression is the most strongly induced by centrosome amplification. Fold induction for this group of genes from RNAseq of MCF10A cells with centrosome amplification relative to controls. (c) Induction of ANGPTL4 expression in cells with centrosome amplification. ANGPTL4 mRNA fold changes from RNAseq in RPE-1 cells in PLK4 versus 608, tetraploids versus parental diploids and tetraploids versus evolved tetraploids. (d) Induction of ANGPTL4 secretion in cells with centrosome amplification. ELISA analysis of secreted ANGPTL4 from cells with or without centrosome amplification. (e) Inhibition of ANGPTL4 compromises paracrine invasion stimulated by cells with centrosome amplification. Shown is the fold increase of MDA-MB468 cells in the bottom chamber of the transwell in the presence of either control IgG or an ANGPTL4 blocking antibody, co-cultured with the indicated MCF10A cells. (f) CRISPR-mediated gene disruption of ANGPTL4 in cells with centrosome amplification inhibits induction of paracrine invasion. Fold increase of MDA-MB468 cells that crossed the Matrigel-coated transwell filter upon co-cultured with control or sgANGPTL4 MCF10A cells with centrosome amplification. All data are means + s.e.m. from n=3 independent experiments, *P < 0.05; analysed with Student’s t - test. **P <0.01, ****P <0.0001; adjusted P value (grey) analysed with DESeq2 Wald test.

### Centrosome amplification activates HIF-1α induction of ANGPTL4

We next addressed the mechanism by which centrosome amplification induces the expression of ANGPTL4. ANGPTL4 is an established target of HIF1 α, the master regulator of the hypoxia response^37, 39, 40^. Thus, we considered the possibility that HIF-1 α is activated in the centrosome amplification induced SASP. HIF1α are crucial transcription factors that regulate oxygen sensing and mediate the hypoxic response^41^. Activation of HIF1α triggers the transcription of genes crucially involved in cancer^42^ and immunity^43^, including cytokines such as IL8, IL6, ANGPTL4, VEGF, and PDGF commonly observed in SASP^20, 37, 44–46^. Consistent with this idea, GSEA demonstrated that a hallmark hypoxia gene signature is present in both MCF10A and RPE-1 cells with PLK4-induced centrosome amplification (FDR q=0.028, Figure 3a and FDR q<0.001, Supplementary Fig 2g). Additionally, newly generated tetraploid RPE-1 cells with extra centrosomes also exhibited enrichment of the hypoxia signature as compared with parental diploid cells (FDR q=0.010, Fig 3b) or evolved tetraploids lacking extra centrosomes (FDR q<0.001, Fig 3c).

**Figure 3.**
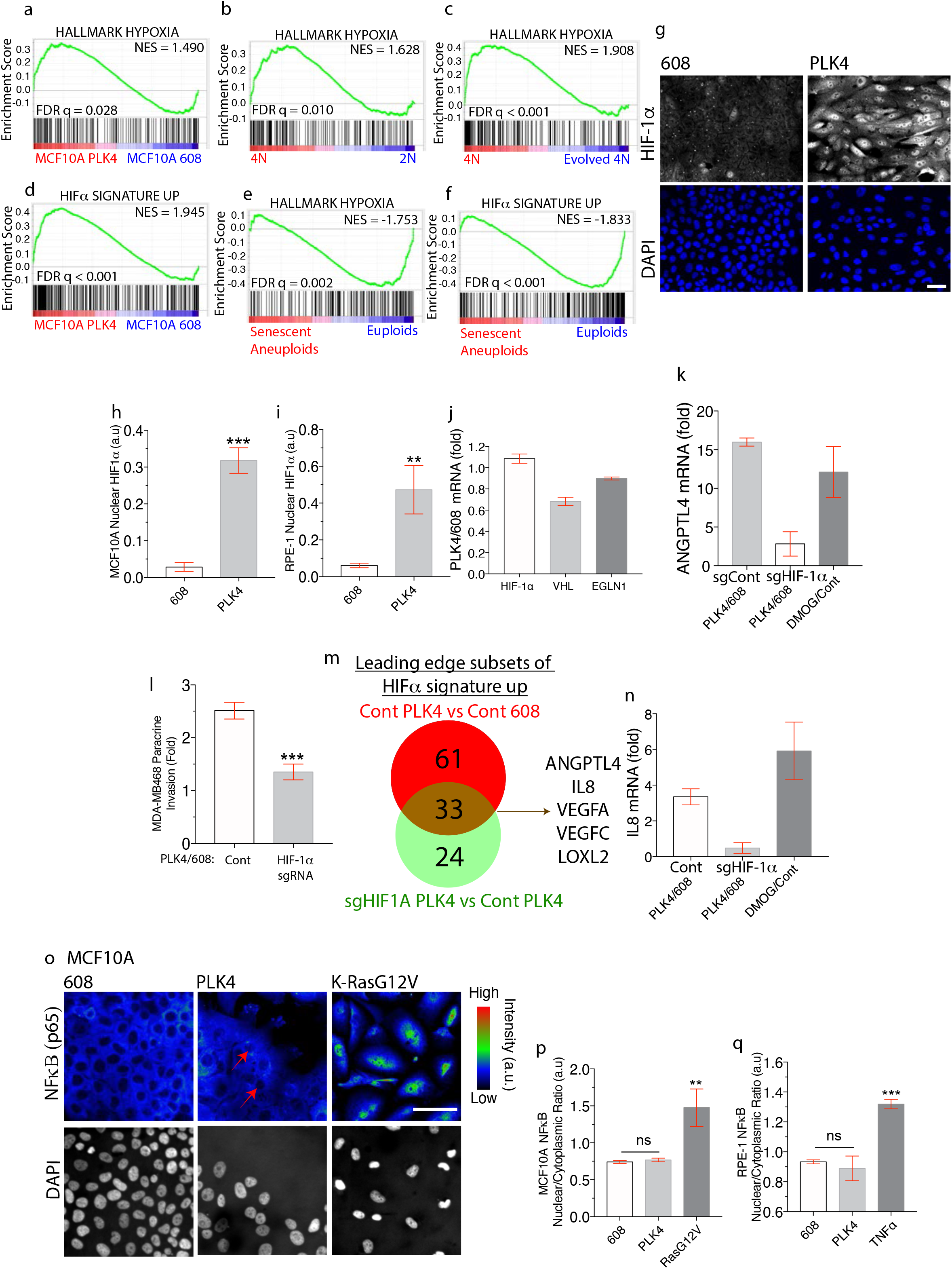
Centrosome amplification activates HIF1α. (a-c) Induction of the expression of hypoxia-inducible genes in cells with centrosome amplification. GSEA revealed strong enrichment of an annotated hypoxia hallmark gene set in cells with centrosome amplification. RNA-seq data sets being compared are indicated at the bottom of the GSEA plots for MCF10A cells (a) and RPE-1 cells (b & c). (d) A custom-generated HIFα signature up gene set in MCF10A cells is strongly induced by centrosome amplification in MCF10A cells. (e,f) Unlike centrosome amplification, aneuploidy-induced senescence suppresses expression of the hallmark hypoxia (e) and custom-generated MCF10A HIFα signature up (f) gene sets. (g-i) Induction of nuclear HIF-1α after centrosome amplification-induced senescence. Representative images (g) and automated quantification of nuclear HIF-1α in MCF10A cells (h) or RPE-1 cells (i) with or without centrosome amplification (see online Methods and Supplementary Fig 4). (j) Centrosome amplification does not alter the expression of HIF-1α, VHL and EGLN1. Fold difference of HIF-1α, VHL and EGLN1 expression levels from MCF10A cells RNA-seq data sets with or without centrosome amplification. (k) CRISPR-mediated gene disruption of HIF-1 α reduces ANGPTL4 expression induced by centrosome amplification. The relative abundance of ANGPTL4 mRNA from RNAseq in centrosome amplification or control hypoxia mimetic DMOG-treated cells is expressed as the fold-change with respect to control cells. Centrosome amplification and control 608 cells are subjected to CRISPR-mediated gene disruption of HIF-1α and compared to CRISPR non-targeting sgRNA controls. (l) CRISPR-mediated gene disruption of HIF-1α in cells with centrosome amplification inhibits the induction of paracrine invasion. Fold increase of MDA-MB468 cells that invaded across the Matrigel-coated transwell when co-cultured with control or sgHIF1α MCF10A cells with centrosome amplification. (m) Forty-nine genes are significantly induced by centrosome amplification induced HIF-1α activation. Venn diagram for the indicated comparisons showing the overlap of the leading-edge subsets of genes from GSEA of HIFα-induced genes. (n) CRISPR-mediated gene disruption of HIF-1α reduces IL8 expression induced by centrosome amplification. Relative abundance of IL8 mRNA from RNAseq in centrosome amplification or DMOG-treated cells is expressed as the fold change relative to controls. Centrosome amplification and control cells are subjected to CRISPR-mediated gene disruption of HIF-1α and compared to the CRISPR non-targeting control. (o) Lack of NF-κB nuclear accumulation after centrosome amplification. Representative images of p65 RELA (NF-κB) and DAPI (grey) in MCF10A cells with and without centrosome amplification as compared with a K-RasG12V positive control. (p,q) Quantification of p65 RELA (NF-κB) nuclear/cytoplasmic ratio in MCF10-A (p) and RPE-1 (q) cells with or without centrosome amplification as compared to K-RasG12V overexpression (p) or TNFα treatment (q). All data are means + s.e.m. from n=3 independent experiments, **P < 0.01, ***P < 0.001,****P <0.0001; analysed with one-way ANOVA, Tukey’s multiple comparison test. Scale bars, 50 μm.

To rigorously define a relevant hypoxia gene set in MCF10A cells, we derived a gene-expression signature of HIF α activation from transcriptomic analysis of MCF10A cells treated with Dimethyloxalylglycine, N-(Methoxyoxoacetyl)-glycine methyl ester (DMOG). DMOG promotes the steady-state accumulation of HIF-1α by inhibiting EGLN/PHD prolyl hydroxlases^47^. Strikingly, this HIFα activation gene set was significantly enriched in both MCF10A and RPE-1 cells after centrosome amplification (FDR q<0.001, Fig 3d and FDR q=0.048, Supplementary Fig. 2h). By contrast, a previously reported data set for aneuploidy-induced senescence in RPE-1 cells^48^ indicated that both the hallmark hypoxia gene set (FDR q=0.002, Fig 3e) and our DMOG-induced gene set (FDR q<0.001, Fig 3f) were suppressed in aneuploid cells, underscoring differences between the effects of centrosome amplification and aneuploidy^18^.

Consistent with these gene expression results, using a HIF-1α specific antibody (Supplementary Fig 2i-k) we find that cells with extra centrosomes exhibited a marked increase in nuclear HIF-1α (Fig. 3g,h,i Supplementary Fig 2l,m). HIF-1α accumulation was post-transcriptional because centrosome amplification did not affect HIF1α mRNA levels (Fig 3j). Moreover, centrosome amplification did not affect the expression of proteins involved in HIF-1α destruction (Fig. 3j). Importantly, CRISPR-mediated gene disruption of HIF-1α in cells with centrosome amplification blocked the activation of HIFα-regulated genes (FDR q=0.014, Supplementary Fig 2n,o), attenuated the induction of ANGPTL4 (Fig 3k), and reduced the capacity of these cells to promote paracrine migration (Fig 3l and Supplementary Fig 2p). A total of 33 genes induced by centrosome amplification were critically dependent on HIF-1 α , including well established HIF-1 α targets such as VEGFA, VEGFC, ANGPTL4, LOXL2, and IL8 (Fig 3m and Supplementary Table 1). Among these, IL8 is an important cytokine that is known to be regulated by HIF-1α^49^. Accordingly, HIF-1α disruption in cells with centrosome amplification reduced IL8 induction (Fig 3n).

The induction of HIF-1α by centrosome amplification occurred regardless of whether cells were cultured in ambient oxygen tension (21% O_2_) or under conditions of physiological normoxia/atmospheric hypoxia (5 % O_2_). Gene expression profiles of RPE-1 cells with or without centrosome amplification were obtained after ~7 days in culture in 5% O_2_. Consistent with our previous results obtained in ambient oxygen, cells in 5% oxygen with centrosome amplification exhibited strong upregulation of gene sets annotated for “extracellular proteins” (FDR q<0.001, Supplementary Fig. 3a), “senescence” (FDR q<0.001, FDR q=0.035, Supplementary Fig. 3b,c,) “hypoxia” and “HIF α activation” (FDR q<0.001, FDR q=0.004, Supplementary Fig 3d,e). Moreover, in physiological normoxia, we observed an enlarged adherent cell size, an ~ 5 fold increase of nuclear HIF-1 α (Supplementary Fig 3f,g), and induction of ANGPTL4 and p21 expression after centrosome amplification (Supplementary Fig 3h).

The activation of HIF-1α that we have identified here is a non-canonical contributor of the SASP secretome. Instead, NF-κB is generally thought to be the primary determinant of the SASP secretome^2^, and DNA damage signalling was shown to activate NF-κB through the transcription factor GATA4^2, 31^. However, centrosome amplification does not induce obvious DNA damage (Fig. 1t,u). Thus, quantitative imaging indicated that centrosome amplification in three cell lines did not result in any detectable increase in the nuclear localisation of NF-κB (Fig. 3o-q and Supplementary Fig. 3i). Because secreted factors regulated by NFκB reinforce senescence^2^, poor NFκB activation likely contributes to the weaker induction of senescence-associated beta-galactosidase activity by centrosome amplification (Fig 1 r,s). Thus, centrosome amplification induces a variant SASP that lacks NF-κB activation. Although evidence suggests that hypoxia and HIF-1α activation can prevent or delay cells from entering senescence^50^. Nevertheless, the HIF-1α activity of cells once they have entered senescence have not been studied. Our data show that centrosome amplification induced SASP constitutes the activation of HIF1α.

### Centrosome amplification activates HIF-1α through induction of ROS

Reactive oxygen species (ROS) is induced by inflammatory cytokines, a common feature of senescence and an established trigger for HIF-1α activation^47, 51^. Therefore, we consider that increased levels of ROS could provide a link between centrosome amplification and HIF1 α activation. Centrosome amplification can activate the Rac1 GTPase^18^. Among its many functions, Rac promotes ROS production because it is a stoichiometric subunit of the NADPH oxidase^34, 52^. Therefore, we tested the hypothesis that after centrosome amplification, the Rac -NADPH oxidase generates ROS, to activate HIF1α. Consistent with this hypothesis, pharmacological inhibition of Rac (Fig 4a and Supplementary Fig 4a), CRISPR-mediated gene disruption of the Rac GTPase exchange factor, Trio^53–55^(Fig 4b, c and Supplementary Fig 4b,c), or siRNA knockdown of a key component of the NADPH oxidase, p22^phox^, (Fig 4d and Supplementary Fig 4d,e,f) all significantly reduced HIF-1α nuclear accumulation in cells with centrosome amplification. By RNA sequencing, CRISPR-mediated gene disruption of Trio also suppressed gene expression signatures associated with HIFα activation, including the induction of ANGPTL4, without significantly affecting HIFα regulated genes in control cells (FDR q=0.0294, Fig 4e, f and FDR q=0.044, Supplementary Fig 4g,h).

**Figure 4.**
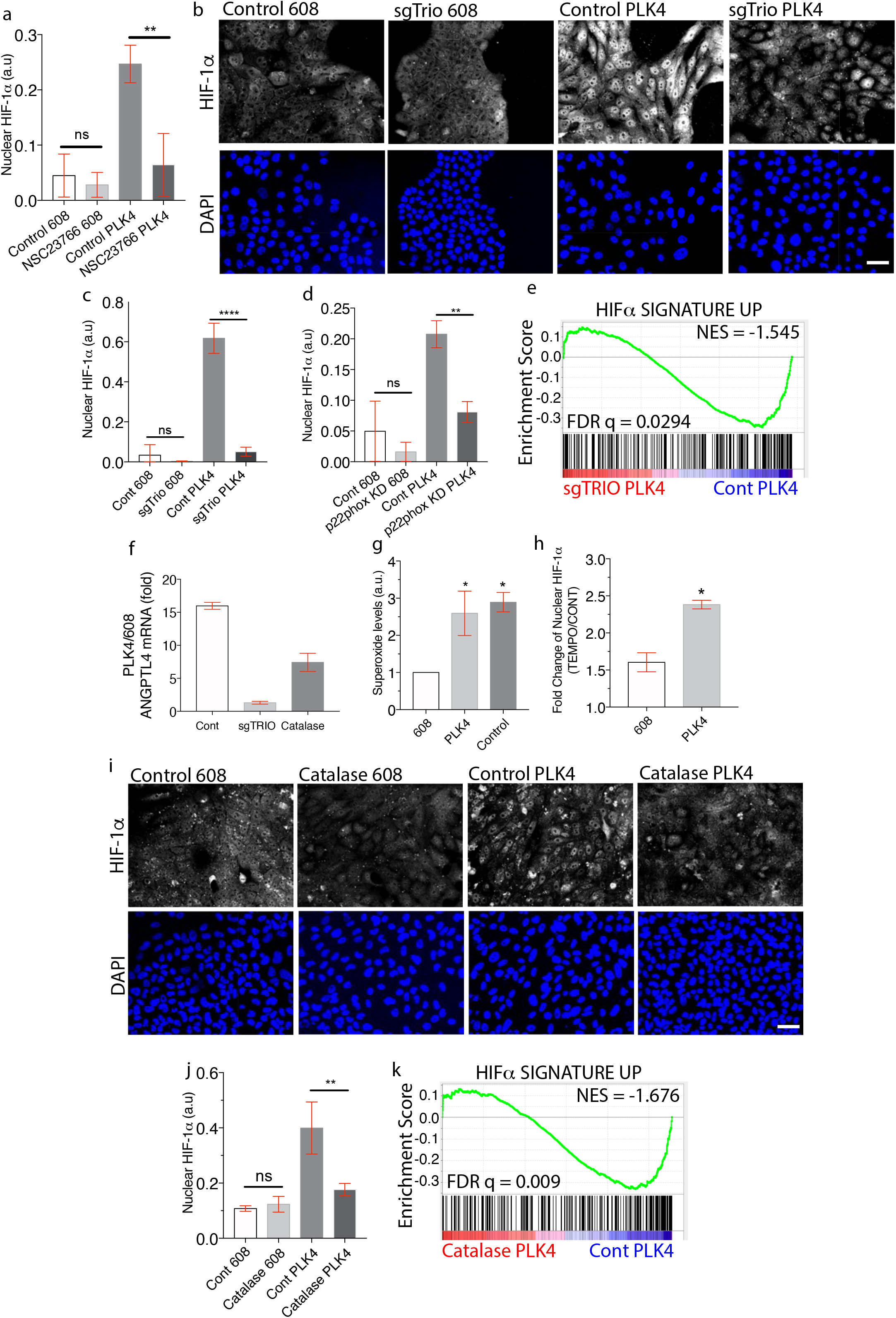
Centrosome amplification induced ROS. (a) Small molecule Rac-1 inhibition prevents the accumulation of nuclear HIF-1α after centrosome amplification. The indicated MCF10A cells were treated with 50μm NSC23766 or vehicle and HIF-1α nuclear accumulation was measured. (b,c) CRISPR-mediated gene disruption of TRIO blocks nuclear HIF-1α accumulation after centrosome amplification. Gene targeting of a pool of cells was performed in the indicated MCF10A cells prior to the initiation of centrosome amplification. Shown are representative image (b) and quantification (c) of nuclear HIF-1α levels in the indicated cells. (d) siRNA knockdown of p22^phox^ inhibits nuclear HIF-1α accumulation after centrosome amplification as in (c). (e) Trio is required for the upregulation of HIFα-induced genes in MCF10A cells with centrosome amplification. Shown is a GSEA plot comparing cells with centrosome amplification with or without TRIO gene disruption. (f) The induction of ANGPTL4 by centrosome amplification requires Trio and ROS. Shown is the fold-induction of ANGPTL4 from RNAseq after centrosome amplification in MCF10A cells after the indicated treatments. (g) Accumulation of superoxide after centrosome amplification Superoxide levels were measured by dihydroethidium-labelling in MCF10A cells with and without centrosome amplification. Pyocyanin treatment is the positive control. (h) Conversion of superoxide into hydrogen peroxide further induces nuclear HIF-1α accumulation in cells with centrosome amplification. Shown are the fold changes in nuclear HIF-1α in the indicated MCF10A cells after treatment with TEMPOL. (i,j) Catalase blocks the nuclear accumulation of HIF-1α in cells with centrosome amplification. Representative images (i) and quantification (j) of HIF-1α in the indicated MCF10A cells with and without catalase medium addition. (k) GSEA showing that catalase treatment prevents the upregulation of the custom HIFα signature up gene set in MCF10A cells. All data are means + s.e.m. from n=3 independent experiments, *P < 0.05, **P < 0:01, ***P < 0:001,****P <0:0001; analysed with one-way ANOVA, Tukey’s multiple comparison test.Scale bars, 50 μm.

Centrosome amplification induces Rac activation^18, 56^. When Rac is activated, NADPH oxidase complex can generate superoxide^57^. Using dihydroethidium labelling, we found that cells with centrosome amplification indeed exhibit higher levels of superoxide (Fig 4g and Supplementary Fig 5a). Cells convert superoxide into the more stable hydrogen peroxide by dismutation of superoxide^58^. Both cellular superoxide and hydrogen peroxide promote HIF1α stabilization^51, 59^. However, when we accelerated the conversion of superoxide into hydrogen peroxide by treating cells with TEMPOL (4-hydroxy-2,2,6,6-tetramethyl piperidin-1-oxyl) ^59^, nuclear HIF1α levels are further increased (Fig 4h and Supplementary Fig 5b), suggesting that that hydrogen peroxide is a major ROS species inducing HIF-1α stabilisation after centrosome amplification. Next, we used catalase addition to the culture medium to directly deplete hydrogen peroxide^60^. Strikingly, in cells with centrosome amplification, catalase treatment prevented nuclear HIF-1α accumulation and suppressed the HIF α -associated gene expression, including the transcriptional induction of ANGPTL4 (Fig 4f,i,j, FDR q=0.009, 4k, and Supplementary Fig 5c). Thus, suggesting that the Rac-NADPH oxidase complex generated ROS activates HIF1α in cells with extra centrosomes. Although ROS induction of HIF-1α is a prominent feature of the SASP from centrosome amplification, elevated ROS is observed in many forms of senescence^30^.

The hypoxia signalling may not be specific to centrosome amplification induced SASP. Indeed, consistent with previous evidence that gamma irradiation activates HIF-1α^61^, nuclear HIF-1α accumulates in γ-irradiation-induced senescent cells, and we observed enrichment of hypoxia-induced genes and a HIFα activation signature (Supplementary Fig 5d-g). HIF-1α is also known to be activated by Ras signaling^62^. Accordingly, we observed enrichment of hypoxia-induced genes and a HIFα activation signature in published data sets^63^ where senescence is induced by oncogene activation (FDR q<0.001, Supplementary Fig 5h,i). Although, the senescence phenotype exhibits significant variability across different cell lines and with different approaches to induces senescenc^15^. Our data show that the expression of hypoxia-induced genes could still be detected in other forms of senescence as well.

In summary, our findings demonstrate that centrosome amplification promotes a variant senescence-associated secretory phenotype (SASP) that constitutes a pathway activating HIF1α (Supplementary Fig 5j). This variant senescence phenotype lacks detectable DNA damage and downstream NFκB activation. Nevertheless, the SASP is well established to generate diverse phenotypic outcomes: promoting tumour invasion in many contexts^64–67^, but also suppressing tumour growth through cellular proliferation arrest and triggering immune response^48, 68^. Thus, it is tempting to speculate that the context-dependent varied consequences of SASP may provide one cell biological basis, for why centrosome amplification promotes malignancy in some models^10, 18^ but are neutral^69, 70^ or inhibitory^71^ in others.

## Acknowledgements

We gratefully thank S.Godinho, D.Pellman, I. Ghobrial, S. Papathanasiou, A. Spektor, A.J. Holland, W. Johnson, S. Seah, S. Mukherjee, H. Zhang, A. Yap, S. Liu, K. Briggs, H. Nicholson, W. Kaelin and K. Polyak for their unstinting support and advice during this project. A.Spektor for gamma-irradiation experiments, L. Hodgson, M. Janiszewska and H. Zhang for technical advice and reagents, F. Abderazza, N. Flanagan, Y. Wang, D. Deconti, and B. Lawney from the Centre for Cancer Computational Biology and X. Qiu and H. Long from Center for Functional Cancer Epigenetics for help with bioinformatics analysis. This research was supported by a Leukemia and Lymphoma Society Fellow Award, USA (5454-17), a Research Scholarship Block Research Fellow Scheme by the Singapore Ministry of Education (C141000207532) and a Young Individual Research Grant (MOH-OFYIRG20nov-0019) by the National Medical Research Council of Singapore to SKW. RP was supported by a Susan G. Komen Postdoctoral Fellowship (PDF15333560). We acknowledge the Nikon Imaging Center at Harvard Medical School for imaging support.

---

**Supplementary Figure 1.**
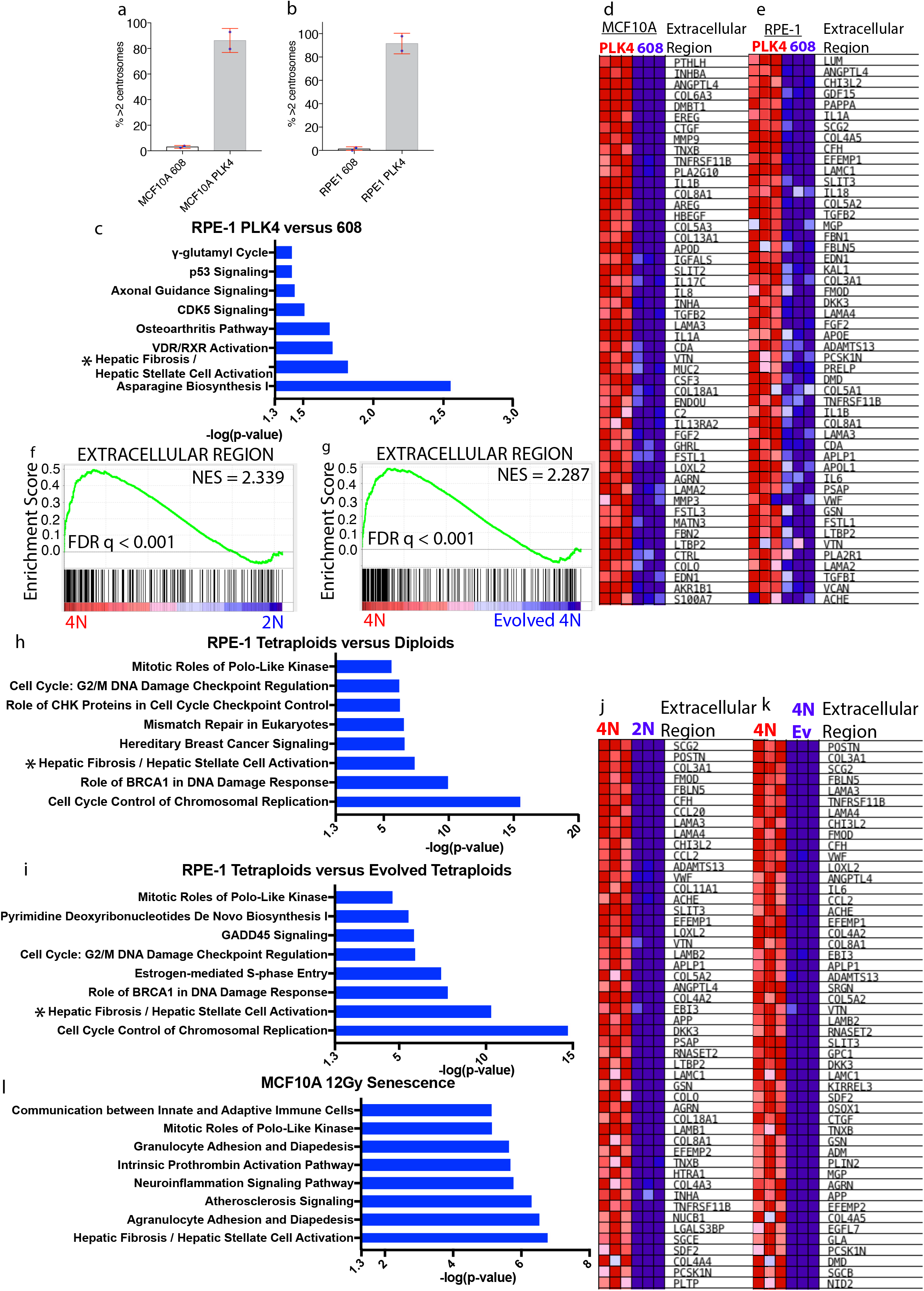
Features of the centrosome amplification SASP. (a,b) The fraction of control (608) and PLK4 induced MCF10A(a) or RPE-1(b) cells with more than 2 centrosomes was determined by immunostaining of centrin-1. Data are means + s.e.m. from n=2 independent experiments. (c) Centrosome amplification alters expression of genes related to cell motility and senescence. Ingenuity Pathway Analysis (IPA) of gene expression changes in RPE-1 cells with centrosome amplification relative to controls revealing the top pathways altered by centrosome amplification. Hepatic Fibrosis/ Hepatic Stellate Cell Activation is a SASP regulated process^22^. (d,e) Induction of the expression of secreted proteins in cells with centrosome amplification. Heatmap showing the leading-edge enrichment of the top 50 extracellular protein expression upregulated in MCF10A (d) and RPE-1 (e) cells with centrosome amplification relative to control. (f,g) Induction of secreted protein expression in cells with centrosome amplification. Gene set enrichment analysis (GSEA) revealed enrichment of genes annotated to be the extracellular region in tetraploids relative to either parental diploids (f) or evolved tetraploids (g). NES: normalised enrichment score; FDR: false discovery rate. (h,i) Centrosome amplification alters expression of genes related to senescence. Ingenuity Pathway Analysis (IPA) of gene expression changes in tetraploids relative to either parental diploids (h) or evolved tetraploids (i) revealing the top pathways altered by centrosome amplification. *Hepatic Fibrosis/ Hepatic Stellate Cell Activation is a senescence regulated process^22^. (j,k) Heatmap showing the leading-edge enrichment of the top 50 extracellular protein expression upregulated in tetraploids relative to either parental diploids (i) or evolved tetraploids (j). (l) γ-irradiation alters expression of genes related to DNA-damage induced senescence. Ingenuity Pathway Analysis (IPA) of gene expression changes in MCF10A cells exposed to 12Gy of γ-irradiation relative to controls revealing the top pathways altered by DNA-damage induced senescence.

**Supplementary Figure 2.**
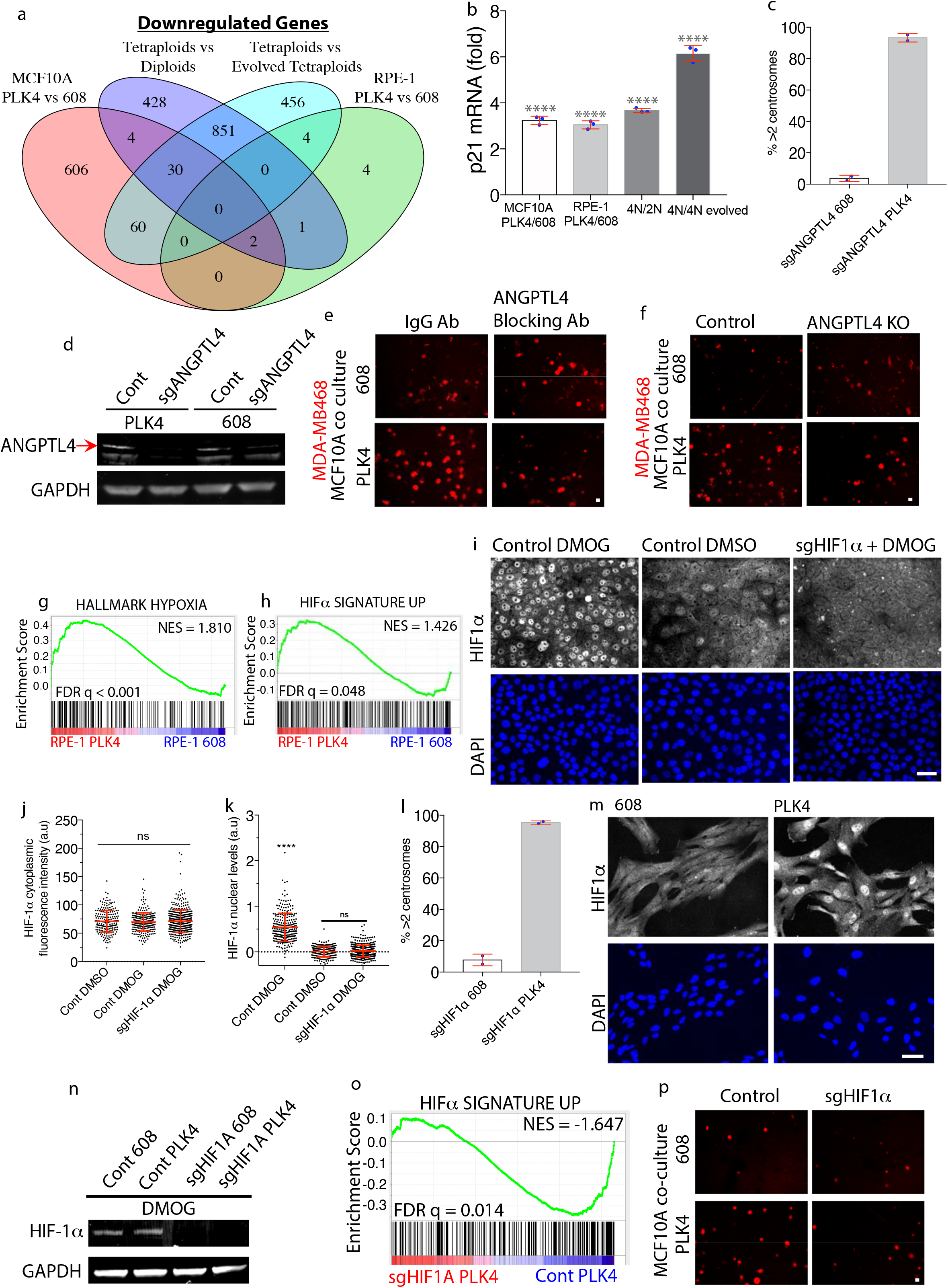
Gene expression altered by centrosome amplification, a functional requirement for ANGPTL4. (a) Zero genes are commonly downregulated by centrosome amplification from all experimental conditions. Venn diagram for the indicated comparisons showing the overlap of down-regulated genes from RNAseq (2 fold change, q<0.05). (b) Induction of p21 expression in cells with centrosome amplification. p21 mRNA fold changes from RNAseq in RPE-1 and MCF10A cells in PLK4 versus 608, tetraploids versus parental diploids and tetraploids versus evolved tetraploids. Data are means + s.e.m. from n=3 independent experiments, ****P <0:0001; adjusted P-value(grey) analysed with DESeq2 Wald test. (c) CRISPR-mediated gene disruption of ANGPTL4 does not affect the induction of extra centrosomes. The fraction of control (608) and PLK4 induced sgANGPTL4 MCF10A cells with more than 2 centrosomes, determined by immunostaining of centrin-1. Data are means + s.e.m. from n=2 independent experiments. (d) CRISPR-mediated gene disruption of ANGPTL4. ANGPTL4 and GAPDH (loading control) immunoblots of lysates from doxycycline uninduced control PLK4, sgANGPTL4 PLK4, control 608 and sgANGPTL4 608 cells. (e) Inhibition of ANGPTL4 compromises paracrine invasion stimulated by cells with centrosome amplification. Shown is the representative image of MDA-MB468 cells in the bottom chamber of the transwell in the presence of either control IgG or an ANGPTL4 blocking antibody, co-cultured with the indicated MCF10A cells. (f) CRISPR-mediated gene disruption of ANGPTL4 in cells with centrosome amplification inhibits paracrine invasion. Shown is the representative image of MDA-MB468 cells that crossed the matrigel-coated transwell upon co-culturing with control or sgANGPTL4 MCF10A cells with centrosome amplification. Scale bars, 50 μm. (g,h) Centrosome amplification upregulates the expression of hypoxia and DMOG-induced genes. GSEA revealed strong enrichment of an annotated hypoxia hallmark gene set (g) and DMOG-induced genes (h) in cells with centrosome amplification. RNA-seq datasets being compared are indicated at the bottom of the GSEA plots for RPE-1 cells. (i) The HIF-1α monoclonal antibody is specific. Representative images of HIF-1α in DMOG-treated, DMSO(vehicle) and DMOG-treated sgHIF-1α cells. (j) HIF-1α cytoplasmic fluorescence intensity is largely background staining. Automated quantification of cytoplasmic mean fluorescence intensity of HIF-1α in DMOG-treated, DMSO(vehicle) and DMOG-treated sgHIF-1α cells. For Cont DMSO, n= 210; Cont DMOG, n=265; sgHIF-1α DMOG, n=464. (k) DMOG induces nuclear HIF-1α. Automated quantification of nuclear HIF-1α above cytoplasmic levels in DMOG-treated, DMSO(vehicle) and DMOG-treated sgHIF-1α cells. For Cont DMSO, n= 210; Cont DMOG, n=265; sgHIF-1α DMOG, n=464. (l) CRISPR-mediated gene disruption of HIF1A does not affect the induction of extra centrosomes. The fraction of control (608) and PLK4 induced sgHIF-1α MCF10A cells with more than 2 centrosomes, determined by immunostaining of centrin-1. Data are means + s.e.m. from n=2 independent experiments. (m) Induction of nuclear HIF-1α after centrosome amplification-induced senescence. Representative images of HIF-1α in RPE-1 cells with or without centrosome amplification. (n) CRISPR-mediated gene disruption of HIF-1α. HIF-1α and GAPDH (loading control) immunoblots of lysates from doxycycline uninduced control PLK4, sgHIF1A PLK4, control 608 and sgHIF1A 608 cells which are DMOG-treated. (o) The custom HIFα signature up gene set is specific for centrosome amplification induced HIF-1α activation. GSEA showing that CRISPR-mediated gene disruption suppresses the custom HIFα signature up gene set in sgHIF1A cells relative to control PLK4 MCF10A cells. (p) CRISPR-mediated gene disruption of HIF1A in cells with centrosome amplification inhibits induction of paracrine invasion. Shown is the representative image of MDA-MB468 cells that crossed the matrigel-coated transwell upon co-cultured with control or sgHIF1A MCF10A cells with centrosome amplification. Scale bars, 50 μ m. All data are means + s.e.m. from 3 independent experiments analysed with one-way ANOVA, Tukey’s multiple comparison test. Scale bars, 50 μm.

**Supplementary Figure 3.**
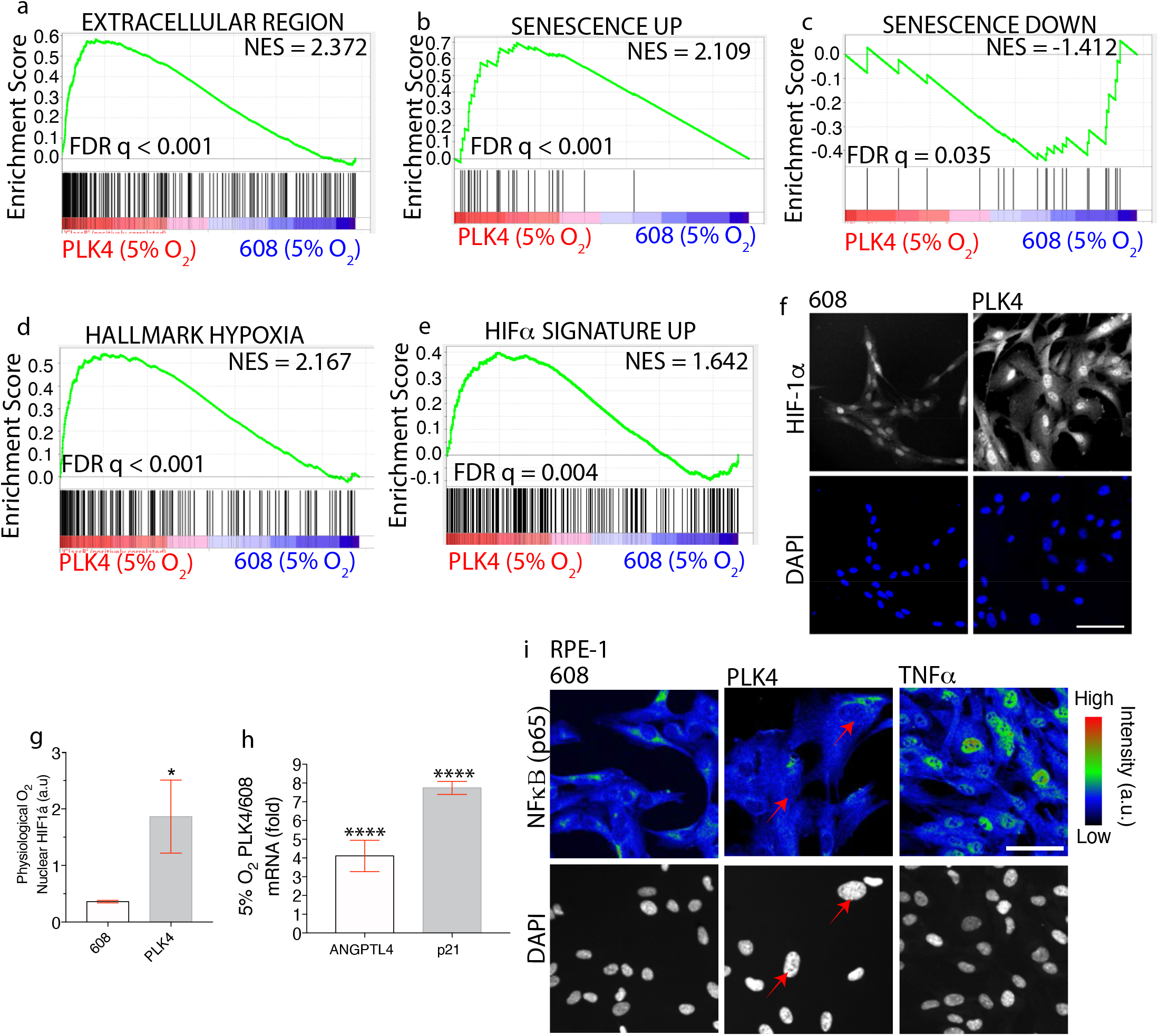
Centrosome amplification induced a SASP that constitutes HIF1α activation independent of prominent NF-κB activity. (a-e) Induction of senescence-associated genes expression in cells with centrosome amplification cultured in physiological normoxia. By GSEA, RPE-1 cells with centrosome amplification cultured in 5% O_2_ induce gene sets associated with the extracellular region(a), genes upregulated(b) and downregulated(c) in senescence, hypoxia(d) and the custom HIFα signature gene set (e). (f,g) Centrosome amplification induces nuclear HIF1α in RPE-1 cells when cells are cultured in 5% O_2_. Representative images (f) and quantification (g) of nuclear HIF-1α in the indicated RPE-1 cells. (h) Fold induction of ANGPTL4 and p21 from RNAseq of cells cultured in 5% O_2_ relative to controls. (i) Lack of NF- κB nuclear accumulation after centrosome amplification. Representative images of p65 RELA (NF- κB) and DAPI (grey) in RPE-1 with and without centrosome amplification as compared with a TNFα treatment positive control.

**Supplementary Figure 4.**
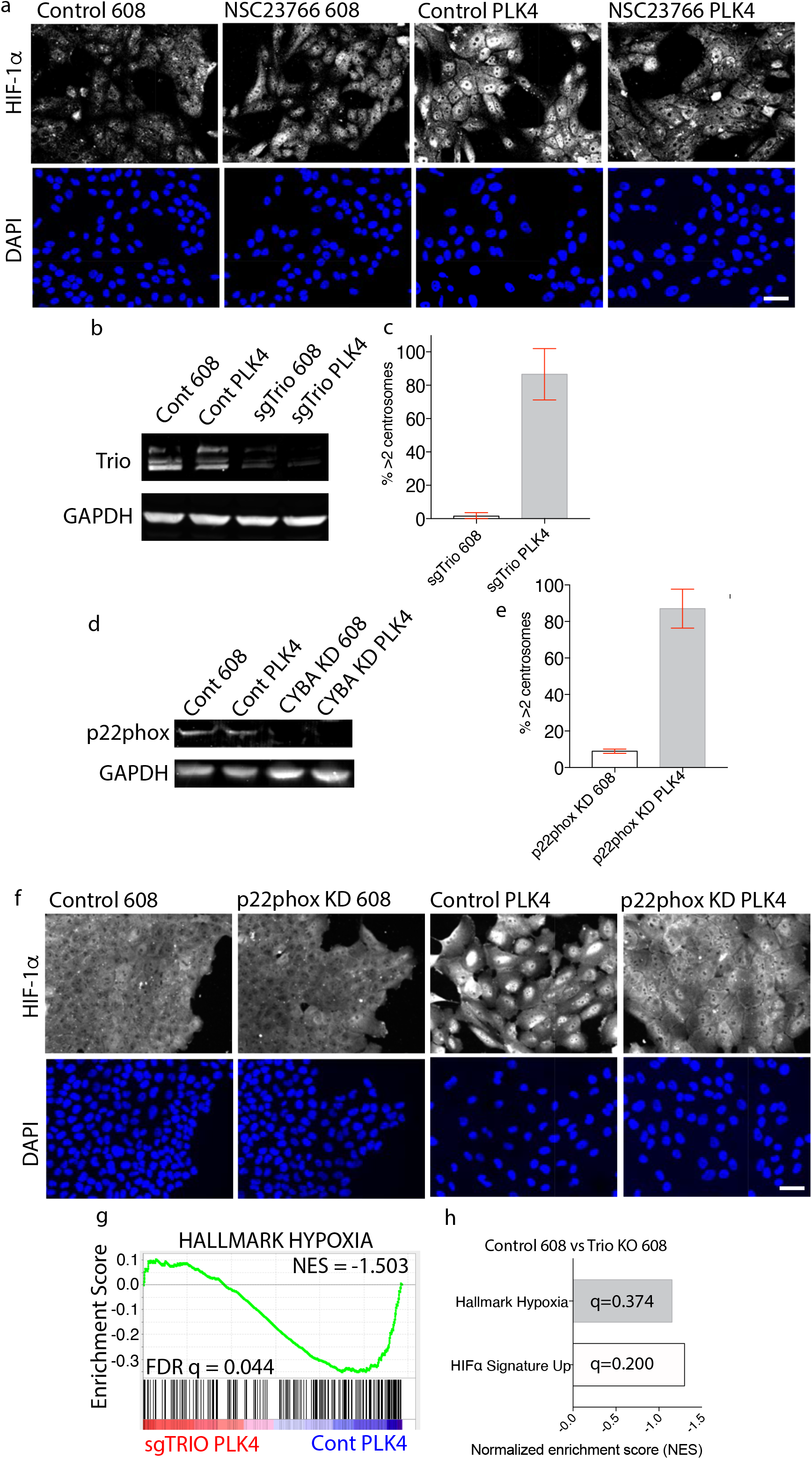
A pathway linking centrosome amplification to HIF-1α activation. (a) Small molecule Rac inhibition prevents the accumulation of nuclear HIF-1α after centrosome amplification. Representative images of the indicated MCF10A cells were treated with 50 μ m NSC23766 or vehicle and HIF-1α nuclear accumulation was measured. (b) CRISPR-mediated gene disruption of TRIO. Trio and GAPDH (loading control) immunoblots of lysates from doxycycline uninduced control PLK4, sgTrio PLK4, control 608 and sgTrio 608 cells. (c) CRISPR-mediated gene disruption of Trio does not affect the induction of extra centrosomes. The fraction of control 608 and PLK4 induced sgTRIO MCF10A cells with more than 2 centrosomes, determined by immunostaining of centrin-1. Data are means + s.e.m. from n=2 independent experiments. (d) siRNA knockdown of p22^phox^. p22^phox^ and GAPDH (loading control) immunoblots of lysates from doxycycline uninduced control PLK4, p22^phox^ knockdown PLK4, control 608 and p22^phox^ knockdown 608 cells. (e) siRNA knockdown of p22^phox^ does not affect the induction of extra centrosomes. The fraction of control (608) and PLK4 induced p22^phox^ knockdown MCF10A cells with more than 2 centrosomes, determined by immunostaining of centrin-1. Data are means + s.e.m. from n=2 independent experiments. (f) siRNA knockdown of p22^phox^ prevents the accumulation of nuclear HIF-1α after centrosome amplification. Representative images of HIF-1α of the indicated MCF10A cells. (g) TRIO is required for the upregulation of hypoxia-induced genes in MCF10A cells with centrosome amplification. Shown is a GSEA plot comparing cells with centrosome amplification with or without TRIO gene disruption. (h) CRISPR-mediated gene disruption of Trio does not significantly affect the regulation of HIF-1α and hypoxia-induced genes in MCF10A control 608 cells. Shown are the normalised enrichment score and false discovery rate q-values of HIF-1α and hypoxia-induced genes for sgTrio 608 cells relative to control 608 cells. Scale bars, 50 μm.

**Supplementary Figure 5.**
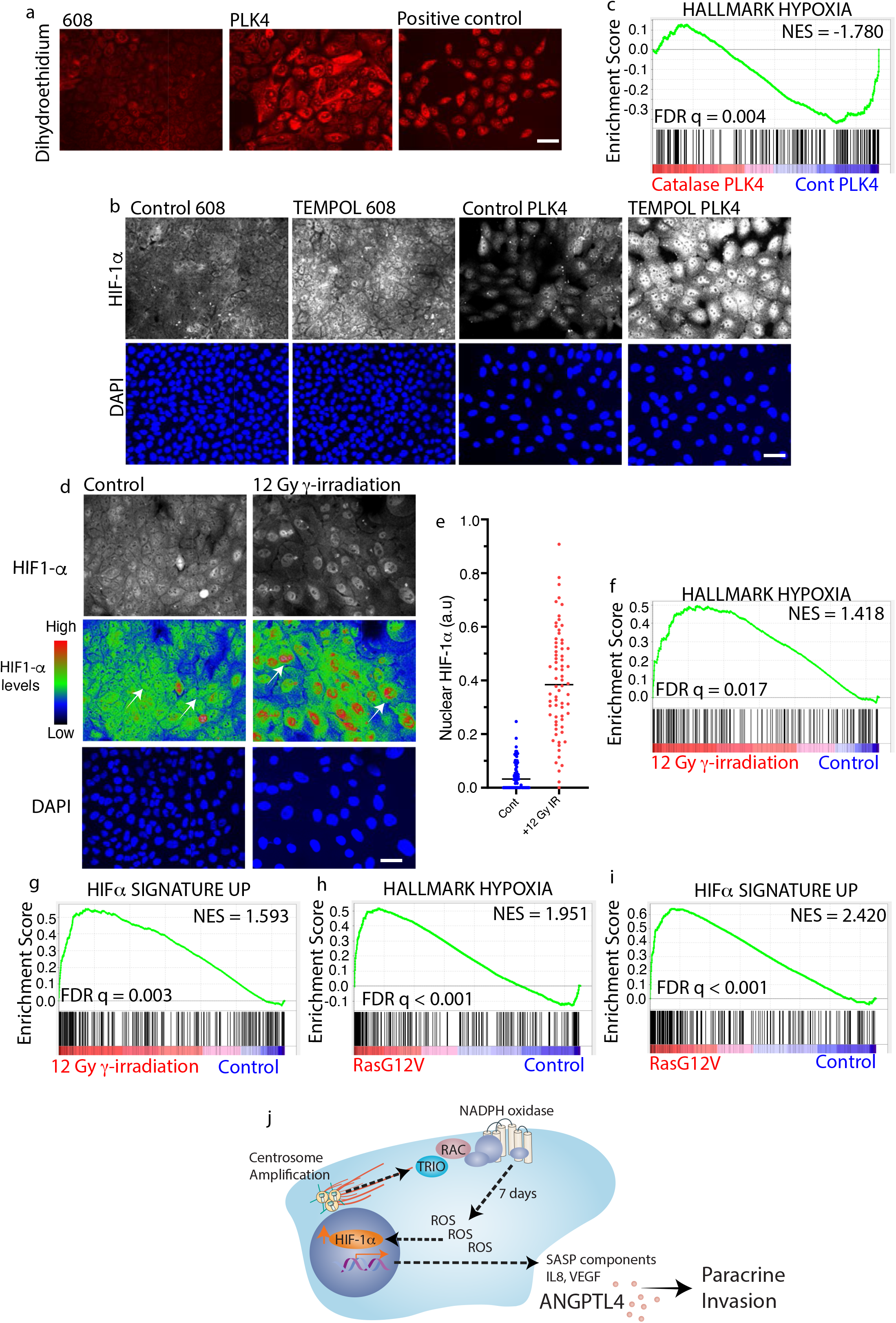
The role of hydrogen peroxide in the activation of HIF-1α after centrosome amplification and activation of HIFα by gamma-irradiation and oncogene-induced senescence. (a) Accumulation of superoxide after centrosome amplification. Representative images of dihydroethidium-labelling of superoxide levels in MCF10A cells with and without centrosome amplification. Pyocyanin treatment is the positive control. (b) Conversion of superoxide into hydrogen peroxide further induces nuclear HIF1α accumulation in cells with centrosome amplification. Representative images of HIF-1α in the indicated MCF10A cells after treatment with TEMPOL. (c) GSEA showing that catalase treatment prevents the upregulation of the hallmark hypoxia gene set in MCF10A cells. (d,e) Nuclear HIF1α levels are significantly increased in senescent cells 8 days post γ-irradiation. Representative images (d) and quantification (e) of nuclear HIF1α (grey) in control and senescent MCF10A cells. (f-i) GSEA showing that the hallmark hypoxia gene sets and custom HIFα signature up (HIFα) are upregulated in MCF10A cells 8 days post γ-irradiation (f,g) and IMR90 cells undergoing oncogene-induced senescence^63^ (h,i). Scale bars, 50 μm. Data are means + s.e.m. from n=3 independent experiments, *P < 0.05; analysed with Student’s t-test. Scale bar, 50 μm.

## METHODS

### Cell culture

All source cell culture was maintained in media containing plasmocin (1:5000 dilution, Invivogen) to prevent mycoplasma contamination. Culture plates were briefly coated with 40μg/ml of fibronectin (Sigma) and glass coverslips were incubated in 40 μg/ml fibronectin for at least 2 hours. Human mammary epithelial MCF10A cells were maintained at 37°C with 5% CO_2_ atmosphere and cultured as previously described^18^. In brief, MCF10A cells were cultured in phenol red-free DMEM/F12 (Invitrogen) supplemented with 5% horse serum (Sigma), 20ng/ml epidermal growth factor (EGF; Sigma), 10mg/ml insulin (Invitrogen), 100 mg/ml hydrocortisone (Sigma), 1ng/ml cholera toxin (Sigma) and 100 IU/ml penicillin and 100 μg/ml streptomycin (Invitrogen). Telomerase-immortalized RPE-1 cells (ATCC), and all derivative cell lines generated in this study were cultured in phenol red-free DMEM/F12 media containing 10% FBS, 100U/ml penicillin, and 100μg/ml streptomycin. RPE-1 cells were maintained at 37°C in 5% CO_2_ atmosphere. For experiments performed in physiological normoxia, RPE-1 cells were maintained at 5% O_2_ conditions in a custom-designed hypoxia chamber (Coy Labs).

To conditionally overexpress PLK4, lentiviral vectors pLenti-CMV-TetR-Blast (17492, Addgene) and pLenti-CMV/TO-Neo-Dest (17292, Addgene) were used. Wild-type PLK4 and PLK4 1–608 cDNAs (control 608) were cloned using the Gateway system into the pLenti-CMV/TO-Neo-Dest vector Cells were coinfected with a lentivirus containing the TetR and with the lentivirus containing the wild-type PLK4 or control 608 open reading frames. Transduced cells were selected with Blasticidin (10 μg/ml) and Geneticin (200μg/ml). control 608 and PLK4 overexpression was induced transiently. Transduced cells that survive the selection are used for subsequent experiments and a maintenance dose of 5 μg/ml of Blasticidin and 80μg/ml of Geneticin was used in stock cultures.

### Generation of tetraploids and ‘’’evolved’ tetraploids with normal centrosome numbers

To generate G1 arrested tetraploids, 15cm dishes were seeded with 6 million exponentially growing RPE1-FUCCI cells, such that they were □65% confluent the following day. Day 2: 4 μM Dihydrocytochalasin B (DCB) was added to each 15 cm dish for 16 hr. Day 3: DCB-treated cells were washed 5 times for 5 minutes, incubated in medium containing 2.5 μg/ml Hoechst dye for 1 hr, then trypsinised in 0.05% trypsin. G1 diploids (2C DNA content; mCherry+, GFP−) and G1 tetraploids (4C DNA content; mCherry+, GFP−) were isolated by FACS sorting. FACS sorted cells were seeded on 6 well plates coated with 40μg/ml fibronectin.

Evolved tetraploid RPE-1 cells, kindly provided by Huadi Zhang, with the normal number of centrosomes were derived as described^26^. Briefly, tetraploid RPE-1 cells were treated for 16 hr with 0.2 μm cytochalasin D, and FACS-sorted by DNA content using Hoechst staining (Molecular Probes). The peak of cells with a DNA content of 8C were isolated and cultured for □1 week before a second iteration of the FACS sorting to re-isolate 8C cells. A total of 5 purifying sorts for □6 weeks was required to generate a pure population of proliferating tetraploid cells with normal centrosome numbers.

### Induction of centrosome amplification and senescence

At day 0, control 608 and PLK4 cells were gently dissociated into single cells, then seeded at low density into media with 2μg/ml of doxycycline such that they were ~80% confluent on day 5. After ~48 hours of doxycycline incubation, the culture media with doxycycline was replaced. On day 5, cells are gently dissociated into single cells before reseeding into 6 well plates to ~80% confluent on day 7. Because control 608 cells grow faster than PLK4 induced cells, control 608 cells are seeded at ~2.5 times lower density than PLK4 induced cells. Cells at ~80% confluency are analysed either at day 7 or day 8. Media is left unchanged for ~3 days before analysis.

### Antibodies, siRNA and drugs

Primary antibodies used in this study were: mouse monoclonal antibodies against Centrin (1:500 for immunofluorescence; Merck Millipore catalogue number 04-1624, clone number 20H5), mouse monoclonal antibodies against ANGPTL4 (1:50 for western blotting; Santa Cruz catalogue number sc-373762, clone number C-7), rabbit polyclonal antibodies against Trio, was a gift from A.Debant (University of Montpellier, France, 1:500 for western blotting), mouse monoclonal antibodies against Rb (Thermo Fisher Scientific catalogue number: BDB554136, clone number G3-245), rabbit polyclonal antibodies against phosphoRb Ser780 (catalogue number 9307S), mouse monoclonal antibodies against GAPDH (1:10000 for western blotting; Abcam catalogue number ab8245), rabbit polyclonal antibodies against GAPDH (1:10000 for western blotting; Abcam catalogue number ab9485), mouse monoclonal antibodies HIF-1α (1:100 for western blotting; BD Transduction Laboratories catalogue number 610959, clone number 54) mouse monoclonal antibodies against HIF-1α (1:50-1:100 for immunofluorescence; Abcam catalogue number ab16066), rabbit polyclonal antibodies against phosphoH2AX (1:500 for immunofluorescence; EMD Millipore Upstate catalogue number 05-636-I, Clone: JBW301), rabbit polyclonal antibodies against p65/RELA (1:500 for immunofluorescence; Santa Cruz Biotechnology catalogue number sc-372), mouse monoclonal antibodies against CYBA (1:100 for western blotting; Santa Cruz catalogue number sc-271968), Mouse IgG antibody (Abcam, ab37355) was used at 40μg/ml as control for blocking experiments. The ANGPTL4 mouse antibody mAb11F6C4, used at 40μg/ml for blocking experiments, was a gift from Tan Nguan Soon, Andrew (Lee Kong Chian School of Medicine, Nanyang Technological University, Singapore). siRNA against CYBA p22phox (Santa Cruz)

Doxycycline (Sigma Aldrich catalogue number D3072) was used at 2 mg/ml. Catalase (Sigma Aldrich (catalogue number: C9322) was used at 2U/μl for 24 hours. TNFα (Roche Diagnostics catalogue number: 11371843001) was used at 20ng/ml for 2 hours. The following doses of inhibitors were used: 50 μM NSC23766 (EMD Millipore), 4μM of dihydrocytochalasin B (DCB; Sigma), 40 ng/ml Doxorubicin (Sigma).

### Immunofluorescence and microscopy

Cells were fixed at 4°C with 4% paraformaldehyde in cytoskeleton stabilisation buffer (10mM PIPES at pH 6.8, 100mM KCl, 300mM sucrose, 2mM EGTA and 2mM MgCl_2_) on ice for 25 min and subsequently permeabilised with 0.2% Triton-X in PBS for 10 minutes at room temperature. Fixed cells were blocked with 5% BSA in TBS overnight at 4°C. Primary antibodies were incubated overnight at 4°C for HIF1α and 45 minutes room temperature for phosphoH2AX and p65/RELA. For centrin-1 staining, cells were fixed with −20°C methanol for 5 minutes on ice. For all immunostaining, secondary Alexa Fluor antibodies were incubated at room temperature for an hour.

Images were collected with Nikon Ti inverted microscope with 20x wide-field optics equipped with perfect focus system, a Hamamatsu ORCA ER cooled CCD camera controlled with Nikon NIS-Element software and an OkoLab 37°C, 5% CO_2_ microscope chamber.

### Matrigel-coated transwell invasion assay

A previously described assay for matrigel-coated transwell invasion assay was adapted for the current study as follows. 8μm Boyden chamber inserts were used (Cell Biolabs: CytoSelect Tumor Transwell ECM Migration Assay, Catalogue No. CBA-216). For the bottom chamber, MCF10A cells with or without centrosome amplification that were cultured for 48 hours in 400μl of full media was used. mCherry MDA-MB-468 cells were seeded on the top chamber. After overnight culture, cells on top of each insert were removed and the trans-well membrane was mounted onto slides. Migration of mCherry MDA-MB-468 cells to the bottom of the membrane was visualised with a spinning disc microscope at 10x magnification.

### CRISPR Knockouts and Lentiviral Transduction

Puromycin selectable CRISPR lentiviruses were obtained commercially from Applied Biological Materials. Three different CRISPR lentiviruses were tested for each target gene. The CRISPR lentivirus that achieves the best knockout and the MCF10A knockout cells were characterised and used for subsequent experiments. The guide RNA targeting the sequences of the genes of interest and scrambled control is as follow. Scrambled control 5’-GCACTCACATCGCTACATCA-3’ catalogue number K011, Trio Target Sequence 1: 5’-?? −3’; catalogue number K2470115, ANGPTL4 Target Sequence 3: 5’-AGGGGCTAACGGGAGGTCGG-3’; catalogue number K0085515

The previously described^72^ HIF-1α pLenti-CRISPRv2 was a gift from William G. Kaelin, Jr. (Dana-Farber Cancer Institute, Boston, USA). The sgRNA sequence targeting HIF1α is as follow: ‘5’-CACCGTGTGAGTTCGCATCTTGATA −’3’.

For lentivirus production, 6μg lentiviral constructs, 3 μg psPAX2 (Addgene), and 3 μg pMD2.G (Addgene) were transfected into 293T cells at 80% confluency in a 10-cm dish with 8 ml of media. 20 μl of lipofectamine 3000 (Life Technologies) was used and transfection was performed according to ‘manufacturer’s instructions. Virus was harvested at 48 hours and 72 hours post-infection, and filtered with a 0.2μm filter. MCF10A cells were infected for 12 hours, with virus containing the genes of interest in the presence of 10 μg/ml polybrene, washed thoroughly, and allowed to recover for 24 hours before any selection.

### Image processing and analysis

ImageJ macros are created to determine the nuclear to cytoplasmic ratio of fluorescence intensity and the number of phosphoH2AX foci in cells. For the quantification of nuclear and cytoplasmic mean fluorescence intensity of HIF-1α and p65, a nuclei mask was generated from the Z-projection of the Hoescht nuclear staining and a 1 μm region surrounding the nucleus was designated as the cytoplasmic region. The watershed function was applied on the nuclei mask to ensure separation of overlapping nuclei.

For the quantification of γ-H2AX foci, a rolling-ball (5 pixels) background subtraction was applied to 8-bit images labelled for γ-H2AX. A nuclear mask generated from the Z-projection of the hoescht nuclear staining after watershed segmentation was used to identify the γ-H2AX foci within an individual nucleus. The ‘Find ‘Maxima’ function on ImageJ was used to count the number of γ-H2AX foci in each nucleus. The noise for the γ-H2AX staining was obtained from cytoplasmic staining and set as the noise tolerance input on the ‘Find ‘Maxima’ function.

### Suspended Cell Size Measurement, SA-β-gal assays and Superoxide Assay

Suspended Cell Size was determined using the coulter counter (Beckman, Vi-cell). SA-β-gal activity was determined using the Senescent Cell Staining Kit (Sigma), by staining SA-β-gal for overnight, according to manufacturer instructions. Cells positive for SA-β-gal were identified by visual scoring. Superoxide levels were measured using a superoxide assay (Abcam catalogue number: ab139476) according to manufacturer instructions.

### NextSeq500 RNA Library Preparation and Sequencing

RNA quantity was determined on the Qubit using the Qubit RNA Assay Kit (Life Tech) and RNA quality was determined on the Bioanalyzer using the RNA Pico Kit (Agilent). Using the NEB Next Ultra RNA Library Prep Kit for Illumina (NEB), which selects for poly(A) mRNA, 100ng of total RNA was used for cDNA library construction, according to the ‘manufacturer’s protocol. Following library construction, DNA quantity was determined using the Qubit High Sensitivity DNA Kit (Life Technologies) and library insert size was determined using the Bioanalyzer High Sensitivity Chip Kit (Agilent). Finally, qPCR was performed using the Universal Library Quantification Kit for Illumina (Kapa Biosystems) and ran on the 7900HT Fast qPCR machine (ABI) for quantification and normalisation of libraries. Libraries that passed quality control were diluted to 2nM DNA content using sterile water and then sequenced on the NextSeq500 (Illumina) at a final concentration of 12pM, according to ‘manufacturer’s instructions.

### RNA Sequencing Analysis

Sequencing reads were aligned to the reference genome using the RNA-specific STAR aligner using default parameters^73^. The quality of the sequencing run and alignment was assessed by RNA-SeQC^74^. The featureCounts tool is used to count sequencing reads mapping to the reference genome at the gene level^75^. The counts are subsequently normalised between samples using DESeq2 to control for experimental variation^76^. Differential expression analysis was performed between the desired experimental contrasts using DESeq2.

### Ingenuity Pathway Analysis

Meta-analysis of the DESeq2 analysed RNA sequencing data was performed to identify the top pathways that are statistically associated with the list of the significantly induced genes^77^ (fold change > 2, FDR q value < 0.05, using the Ingenuity Pathway Analysis platform IPA, Ingenuity Systems, Qiagen).

### Gene Set Enrichment Analysis

We used GSEA v3.0 to examine the association between gene sets and gene expression. Normalised read counts from DESeq2 analysis were used for GSEA. Genes with a 0 read count in either control or experimental group were excluded. GSEA was performed using gene sets as permutation type, 1,000 permutations and log2 ratio of classes as metrics for ranking genes. We employ the hallmark gene sets from the Molecular Signatures Database (MSigDB)^25, 78, 79^. Hallmark gene sets summarise and represent specific well-defined biological states or processes and display coherent expression. These gene sets were generated by a computational methodology based on identifying gene set overlaps and retaining genes that display coordinate expression, thus reducing data noise and redundancy.

To construct a HIFα signature up gene set we performed RNA sequencing of MCF10A cells after overnight treatment with 1μm of DMOG, with DMSO treated cells as control. We included genes that were upregulated by DMOG in DESeq analysis (p(adjusted)<0.05, fold change > 4, Supplementary Table 1).

### Quantitative RT-PCR

Total RNA was isolated using the RNAeasy Mini Kit (Qiagen), and cDNA was synthesised using SuperScript III (Invitrogen) according to manufacturer instructions. Quantitative RT-qPCR was performed in triplicate using SYBR Green (Life Technologies) on a ViiA 7 Real-Time PCR machine (Thermo Fisher Scientific) with GAPDH as an internal normalisation control.

qPCR primers (PrimeTime® Predesigned qPCR Assays) purchased from IDT were used and product information were as follow: GAPDH, Assay ID: Hs.PT.39a.22214836; Angptl4, Assay ID: Mm.PT.58.12348309.

### ELISA

Cells were starved for 2 hours before reintroducing full media to collect cellular secretion. After 24 hours of incubation, the conditioned (full) media is collected and a protease inhibitor cocktail (Sigma) was added to prevent protein degradation. The conditioned media was centrifuged for 10 minutes at 5000rpm to remove any cellular debris and subsequently quantified for ‘ANGPTL4’s concentration with ELISA (ELH-ANGPTL4, RayBiotech). The total cell number was used to normalise ‘ANGPTL4’s concentration across replicates.

## Notes

### Competing Interest Statement

The authors have declared no competing interest.

